# A network regularized linear model to infer spatial expression pattern for single cells

**DOI:** 10.1101/2021.03.07.434296

**Authors:** Chaohao Gu, Zhandong Liu

## Abstract

Spatial gene-expression is a crucial determinant of cell fate and behavior. Recent imaging and sequencing-technology advancements have enabled scientists to develop new tools that use spatial information to measure gene-expression at close to single-cell levels. Yet, while Fluorescence In-situ Hybridization (FISH) can quantify transcript numbers at single-cell resolution, it is limited to a small number of genes. Similarly, slide-seq was designed to measure spatial-expression profiles at the single-cell level but has a relatively low gene-capture rate. And although single-cell RNA-seq enables deep cellular gene-expression profiling, it loses spatial information during sample-collection. These major limitations have stymied these methods’ broader application in the field. To overcome spatio-omics technology’s limitations and better understand spatial patterns at single-cell resolution, we designed a computation algorithm that uses glmSMA to predict cell locations by integrating scRNA-seq data with a spatialomics reference atlas. We treated cell-mapping as a convex optimization problem by minimizing the differences between cellular-expression profiles and location-expression profiles with a L1 regularization and graph Laplacian based L2 regularization to ensure a sparse and smooth mapping. We validated the mapping results by reconstructing spatial-expression patterns of well-known marker genes in complex tissues, like the mouse cerebellum and hippocampus. We used the biological literature to verify that the reconstructed patterns can recapitulate cell-type and anatomy structures. Our work thus far shows that, together, we can use glmSMA to accurately assign single cells to their original reference-atlas locations.

---

Precise gene-expression spatial regulation at the individual cell level is critical to normal development and disease pathogenesis. For example, asynchronous cell-proliferation in Drosophilia depends upon Hippo-signaling-pathway spatial expression during early embryogenesis^3^. Single-cell RNA sequencing (scRNA-seq) allows us to measure many genes for each individual cell. But spatial information is often lost after tissue-digestion and cell-dispersion. We therefore propose a new algorithm—glmSMA—which, by integrating the reference atlas per the Slide-seq or FISH-based method, can accurately map individual cells back to their original locations in complex tissues.

Current spatial-omics techniques include the next-genome-sequencing (NGS)-based method and imaging-based method. Both methods have significant limitations. On the NGS side, slide-seq’s genome-wide measurement allows us to obtain spatially resolved gene-expression data at the single-cell level but has a low gene-capture rate and produces relatively small barcoded beads and, therefore, a limited measurable area^6^. Similarly, spatial transcriptomics like 10x Visium provide high-quality mRNA transcripts in a larger area, but the resolution is much lower than with Slide-seq^12^. On the image-based side, FISH-like methods can provide tissue mRNA levels with high resolution but can only measure a limited number of genes. We tested glmSMA in several tissues based on different spatial-omics techniques including both NGS and imaging-based method, and the cell assignments were still accurate.

Several algorithms have been proposed to spatially identify gene-expression in tissues-of-interest at a single-cell level^1,2,4,9,10^,^19^. However, no one has yet tested whether spatial transcriptomics data or Slide-seq data is a good reference for spatial-location-prediction at a single-cell level. For example, Dist-map used FISH images from the BDTNP fly-embryo database but can only retrieve tens of genes to build a reference atlas^2^. Likewise, NovoSparc cannot integrate information from spatial references^1^; rather, it relies purely on marker-gene-expression while assuming expression-local similarity. It was also mainly tested in the simulated data set. Also, most spatial-mapping algorithms locate one cell-group instead of sorting individual cells into regions. When MIA^9^ was tested in pancreatic tumors using spatial transcriptomics data, for example, it assigned 3 or 4 predefined cell types from ~1,000 cells into several large domains. HMRF^10^ used smFISH mouse-brain images as a reference atlas but mapped fewer than ten cell-types back to their original regions. While both of these methods used scRNA-seq data and the reference atlas, they didn’t directly assign individual cells back to their locations; instead, they simplified the problems by first grouping the cells and then assigning cell-groups to larger areas. There had been no evidence that integrating Slide-seq and scRNA-seq data can map individual cells back to their original locations. Our GlmSMA, however, combined the two platforms to accurately predict cell-assignments.

## GlmSMA accurately reconstructs intestinal villus *in vivo*

To determine whether individual cells in the intestinal villus can be correctly assigned to their original locations, we performed glmSMA on 1,148 cells derived from 6 distinct intestine zones. We found a monotonical relationship between the cell-to-cell physical distance and cell-to-cell expression correlation (Extended Data Fig. 2). Using the top 100 variable genes, we were able to correctly map ˃99% of the cells to within one layer of their original layer (Extended Data Fig. 1). These results indicate that our algorithm accurately assigns cells into one-dimensional tissues.

## GlmSMA successfully assigns single cells back to their original locations in both in silico and *in vivo* Drosophila embryos

We next tested our algorithm on more complicated, two-dimensional tissue from the Drosophila embryo. We first established the reference profile using an expression atlas of 84 marker genes in developmental stage 5 that we retrieved from the Berkeley Drosophila Transcription Network Project (BDTNP)^5^. BDTNP-database embryos are collected at developmental stage 5 and fluorescently stained to label marker-genes’ nuclei and expression patterns (Extended Data Fig. 5).

To evaluate glmSMA’s performance in complex tissues, we simulated scRNA-seq data on 1,000 individual cells from a randomly selected patches in BDTNP reference atlas using a strategy similar to Nitzan et al’s^1^. We then tested whether glmSMA could map these 1,000 simulated scRNAseq profiles back to the 2D embryo’s surface. As with the intestinal villus dataset, we found a monotonically increasing relationship between cell-to-cell physical distance and cell-to-cell expression distance (Fig. 2a). Using 60 marker genes, we correctly mapped ˃99% of the cells back to their original positions. (Fig. 2b) We also tested whether the L1 and generalized L2 regularization induced a sparse solution. Without regularization, our model yielded multiple optimal locations for each cell. With regularization, a sparse model induced far fewer predicted locations resulting in fewer false positives (Fig. 2c). Our results were still robust after we resampled the original images into 1,500 and 500 patches. To evaluate the predicted cell assignments’ accuracy, we compared the reconstructed spatial-expression patterns on marker genes (e.g., *ftz*, *tsh* and *gt*) to these genes’ experimental patterns from virtual FISH (Fig. 2d). We observed that the patterns are highly similar.

**Fig. 1:**
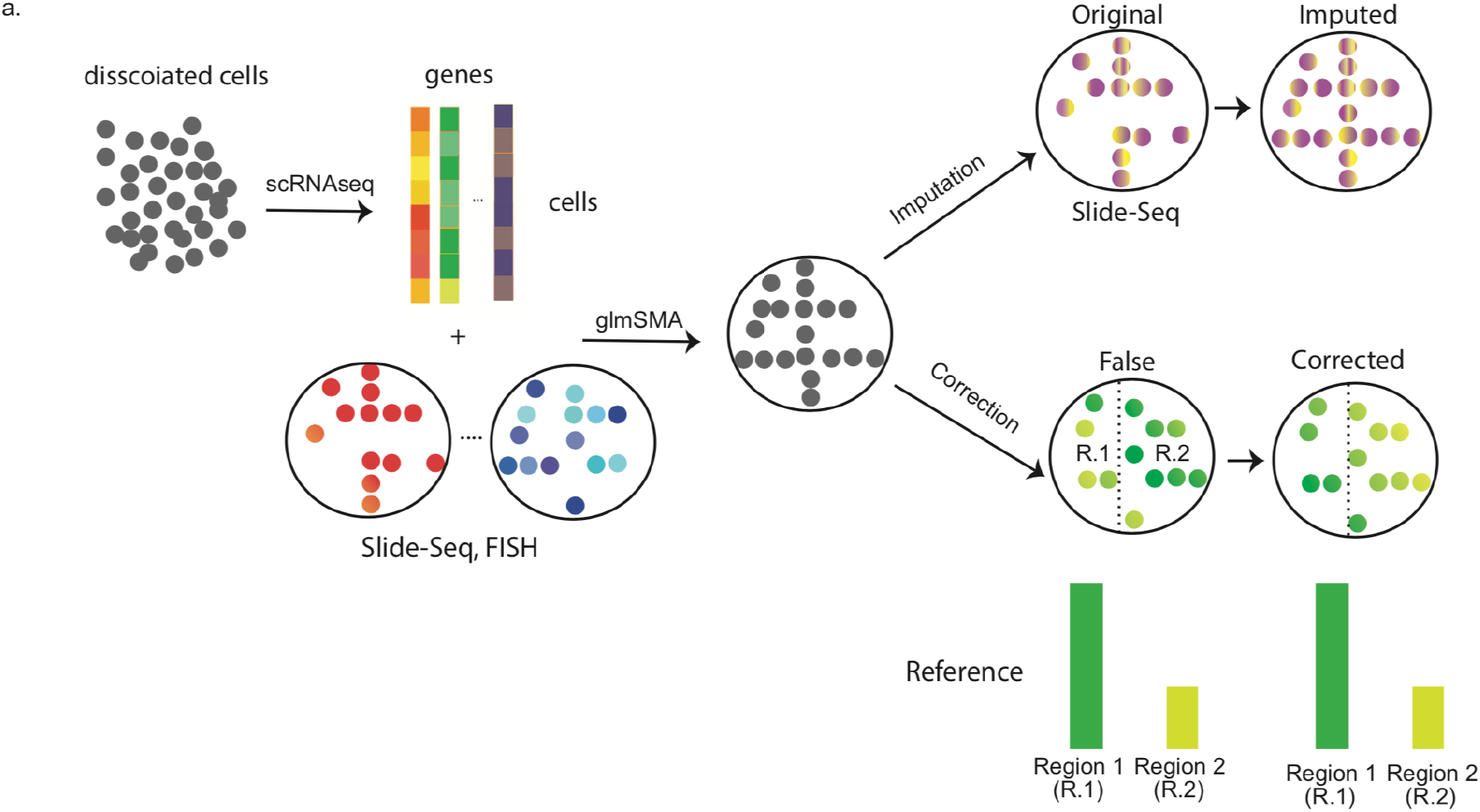
Overview of glmSMA. **a**, The input are the gene expression matrix by scRNA-seq from dissociated cells and the location expression matrix of reference atlas by Slide-seq, FISH, or other spatial-omics technologies. The output is the predicted locations of single cells in the reference atlas. By using the mapped cells and scRNA-seq profiles, we can impute or correct the original patterns which are not captured by the original spatial-omics technologies.

**Fig. 2:**
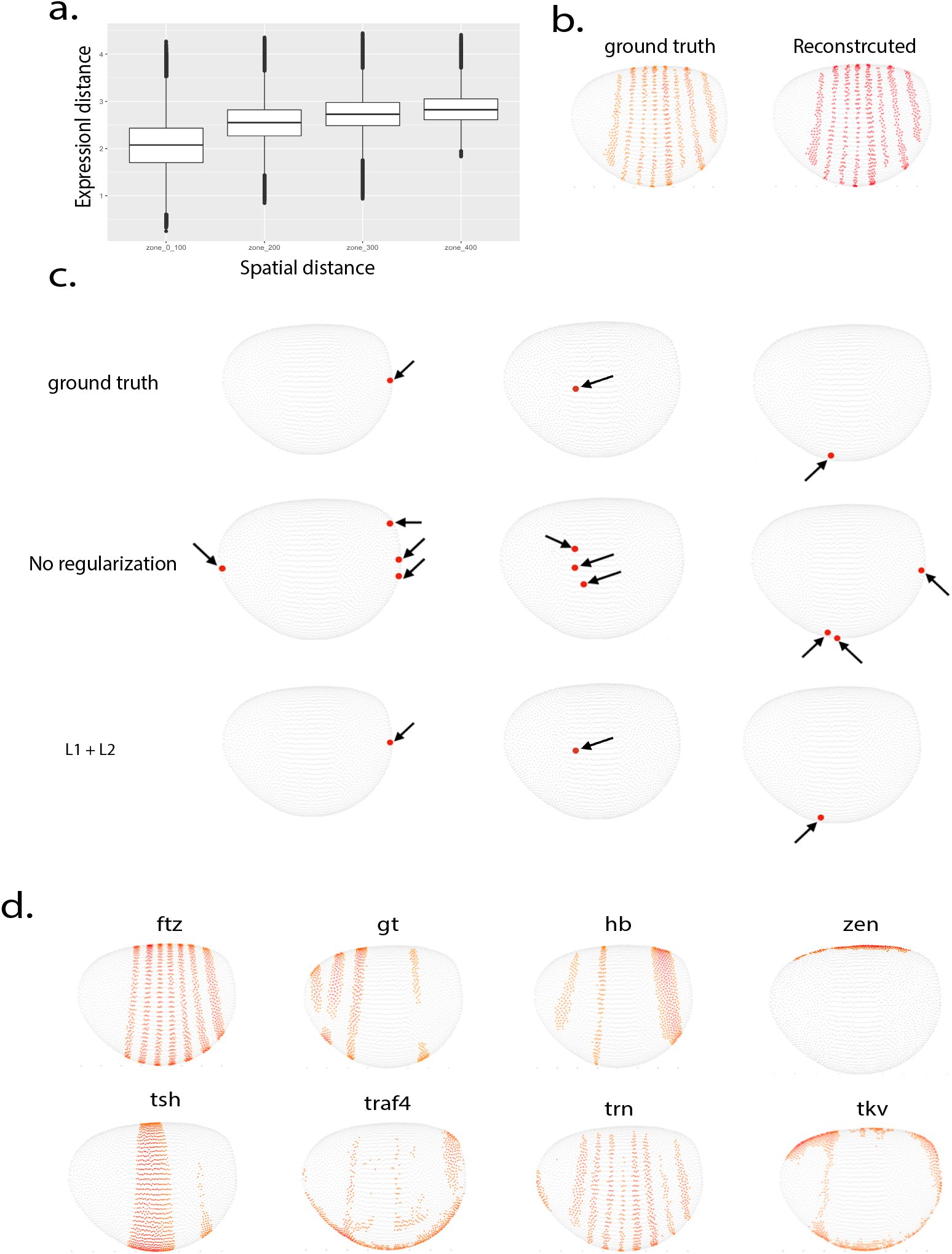
GlmSMA successfully reconstructs the Drosophila embryo on the simulated dataset based on the BDTNP database. **a**, Cell-to-cell physical distance increased monotonically with the cell-to-cell expression distance. **b**, Mapping results of selected cells in the Drosophila embryo. Over 99% of the selected cells were correctly assigned to their original locations. **c**, Regularization path in the cell assignments. Without regularization, cells are assigned to multiple locations. With L1 and generalized L2 regularization, individual cells are more likely to be assigned to one unique position or a very small patch. **d**, Reconstructed spatial expression patterns of marker genes from the mapping results. The reconstructed patterns were consistent with the Virualfly database.

Next, we used a real scRNA-seq dataset to assess glmSMA’s mapping accuracy. We extracted a 1,297-cell drosophila scRNA-seq dataset with approximately 800 genes at developmental stage 5. In the t-SNE plot, the original scRNA-seq study reported four clusters (Fig. 3a). By mapping single cells into a virtual embryo surface, we observed that these four groups actually have distinct spatial locations. Specifically, the black cluster represents cells located in the anterior and posterior. The Blue cluster consists mostly of ventrally located cells while the red and green spots show cells spanning from ventral to dorsal (Fig. 3a). Due to the lack of marker genes and redundancy in the expression patterns, on average, 5 cells were mapped to the same location. We found that these locations are not identifiable because their expression profiles are so similar to those of the 84 marker genes (Extended Data Fig. 15).

**Fig. 3:**
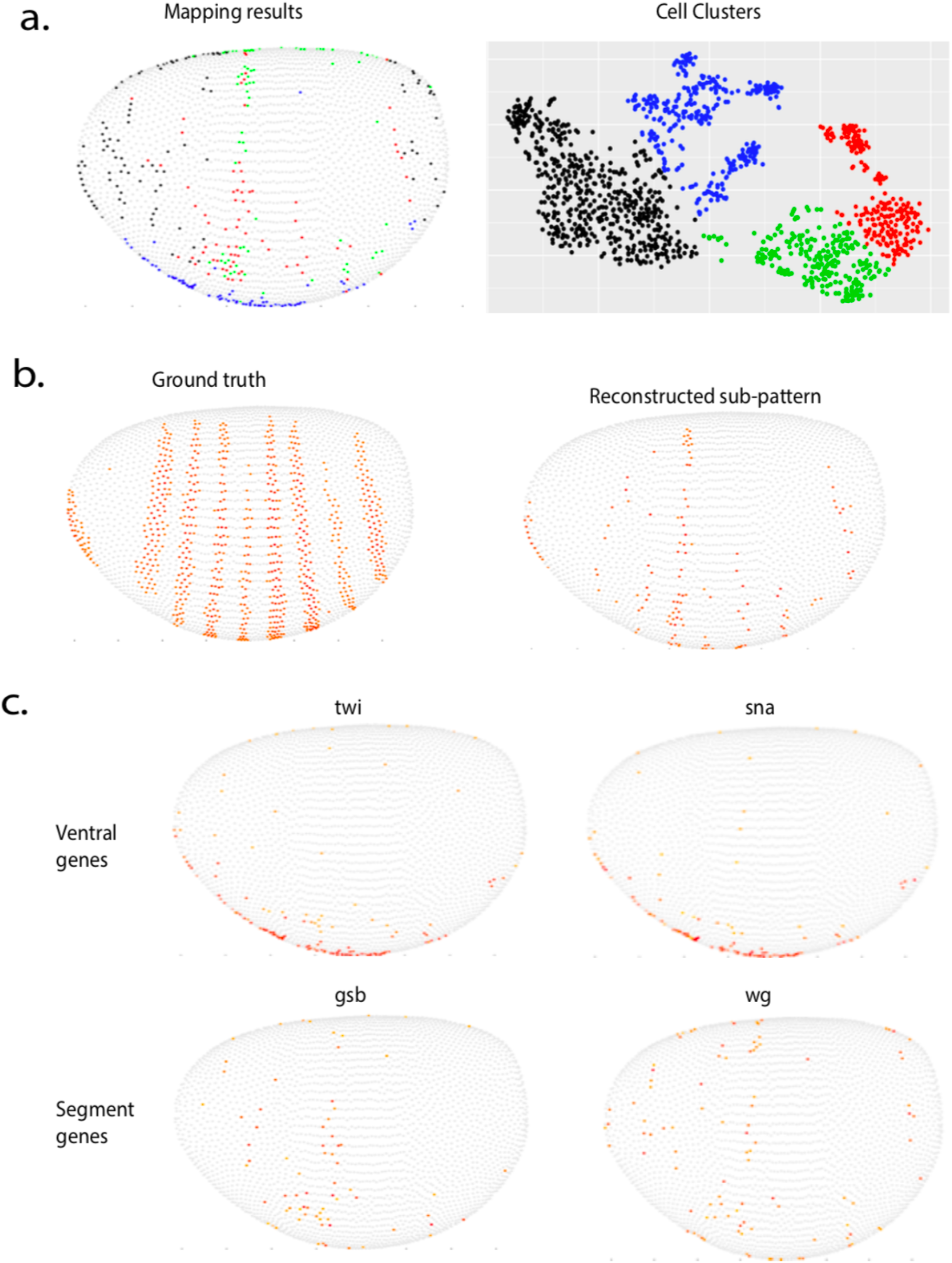
GlmSMA accurately reconstructed the Drosophila embryo from a scRNA-seq dataset. **a**, 1,297 cells were clustered into four clusters. Cells were assigned back to the embryo using glmSMA. Blue cells are mainly clustered in the ventral part; red and green cells are mostly located in the middle part of the embryo; the black cells occupy the anterior and posterior regions. Cells with the same color have closer physical distance. **b**, Example of original and predicted expression patterns for the segment marker genes. GlmSMA successfully reconstructed the partial expression patterns compared to the original one. **c**, Examples of reconstructed spatial expression patterns for the well-known marker genes in the embryo development. *Twi* and *sna* were ventral genes that were highly expressed in the ventral regions. *Gsb* and *wg* were segment genes and their reconstructed patterns were consistent with literature.

To further investigate whether glmSMA accurately predicts spatial gene-expression, we compared the reconstructed important developmental-marker genes’ patterns with virtual FISH patterns. These developmental-marker genes often exhibit special spatial expression patterns (Fig. 3b & Extended Data Fig. 6). For example, *snail* expression at stage 5 has a ventral-to-dorsal gradient (Fig. 3c). We also investigated dorsal- and gap-genes’ spatial-expression patterns (e.g., *twi*, *gsb* and *wg*), and all the reconstructed spatial- expression patterns were consistent with the literature^14^. Most ventral cells were spatially distributed in the ventral area, where *sna* and *twi* were highly expressed, while *gsb* and *wg* were spatially expressed as segmentation stripes (Fig. 3c). We then tested whether L1 and generalized L2 regularization induced a better solution for the cell assignments. With L1 and generalized L2 regularization, cells were assigned into small patches instead of multiple scattered locations (Extended Data Fig. 18). Together, these verification results demonstrate that the mapped cells went to plausible positions.

## GlmSMA successfully locates single cells in the mouse cerebellum both *in silico* and *in vivo*

To further demonstrate that glmSMA can be applied in more complex tissues, we assessed its performance in the mouse cerebellum. We first used a recently developed spatial-omics technology—Slide-seq—to establish the mouse-cerebellum reference atlas (Extended Data Fig. 7,8,9). One sagittal section of the mouse cerebellum in the Slide-seq dataset contained 46,376 locations, and one location had 1−1.5 individual cells with 19,782 genes. We next simulated the scRNA-seq data by using a pipeline similar to Nitzan et al’s to bin neighboring reference-atlas locations and coarse-grain the cerebellum^1^. After simulation, we tested whether glmSMA can map the simulated 7,724 cells back into their original 46,376 locations using the top 100 variable genes. We found that the pairwise physical distance between cells increased monotonically with the pairwise-expression distance between cells (Fig. 4a).

**Fig. 4:**
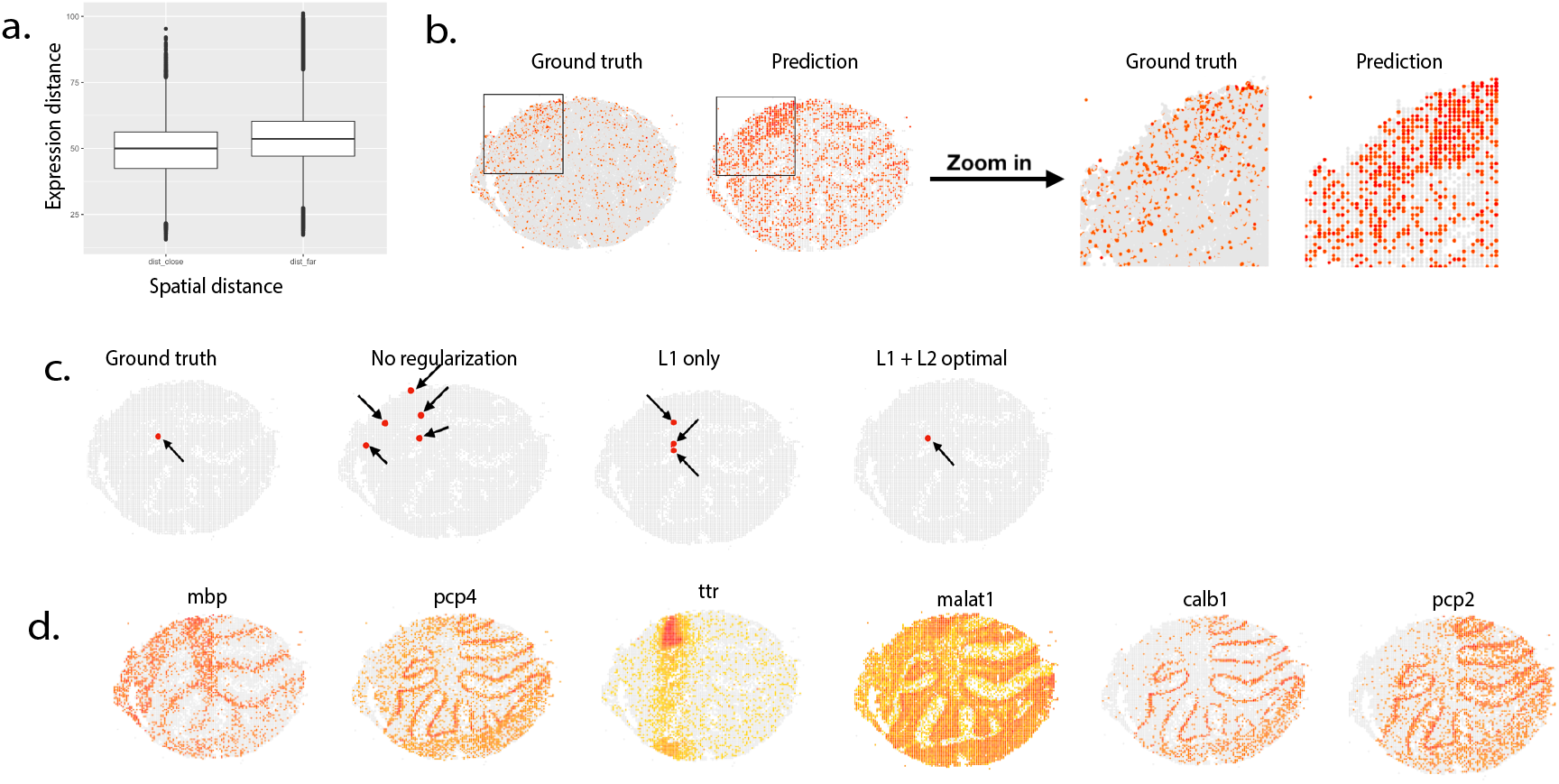
GlmSMA accurately reconstructed the spatial expression patterns in the mouse cerebellum. **a**, Montonocal relationship between cell-to-cell physical distance and cell-to-cell expression distance. **b**, Mapping results of simulated single cells from Slide-seq compared with the ground truth. Over 99% of the cells can be accurately assigned to their original locations. **c**, Regularization path for the cell assignments. By setting lambda1 = 0.51 and lambda2 = 0.32, glmSMA can achieve good performance with L1 and generalized L2 regularization. Cells can be assigned to small patches or unique positions. **d**, Examples of reconstructed spatial expression patterns of well-known marker genes.

As expected, glmSMA correctly assigned ˃99% of the simulated cells to their original locations (Fig. 4b). As we decreased the number of marker genes to 40, glmSMA’s performance dropped slightly, but still assigned ˃97% of cells to the correct locations (Extended Data Fig. 19). When we further tested whether L1 and generalized L2 regularization can generate a better solution for cell assignments, we found that, after regularization, cells can be assigned into either one or a small number of locations instead of multiple scattered ones, resulting in fewer false-positive assignments (Fig. 4c & Extended Data Fig. 14). To further validate the cell assignments, we reconstructed specific marker-genes’ spatial-expression patterns. We demonstrated that the majority of gene patterns recapitulated the original images from the Slide-seq (Fig.4d).

We next used a real biological dataset to assess glmSMA’s ability to locate cells in complex tissues. We retrieved the scRNA-seq data from the online database DropViz^16^ and clustered single cells into 11 cell-types using predefined marker genes, in which most of the cells were classified as granule and purkinje cells by DropViz (Fig. 5b). We randomly selected 1,000 cells and 7,724 positions were available for mapping. Due to the cell locations’ lack of ground truth, we validated the mapping results in two ways. First, we examined the spatial cell-types in the assigned locations on the mouse-cerebellum anatomical map. As expected, most of the granule cells were correctly assigned to the cerebellar cortex, where they were surrounded by the Purkinje cells, and one clear spatial boundary existed between the two cell-types (Fig. 5a). Second, we reconstructed spatial expression patterns of well-known marker genes that were highly expressed in specific cell-types (Extended Data Fig. 20). For example, prior research shows that tulp4 was highly expressed in the Purkinje cells^16^; our reconstructed patterns were consistent with this (Fig. 5b).

**Fig. 5:**
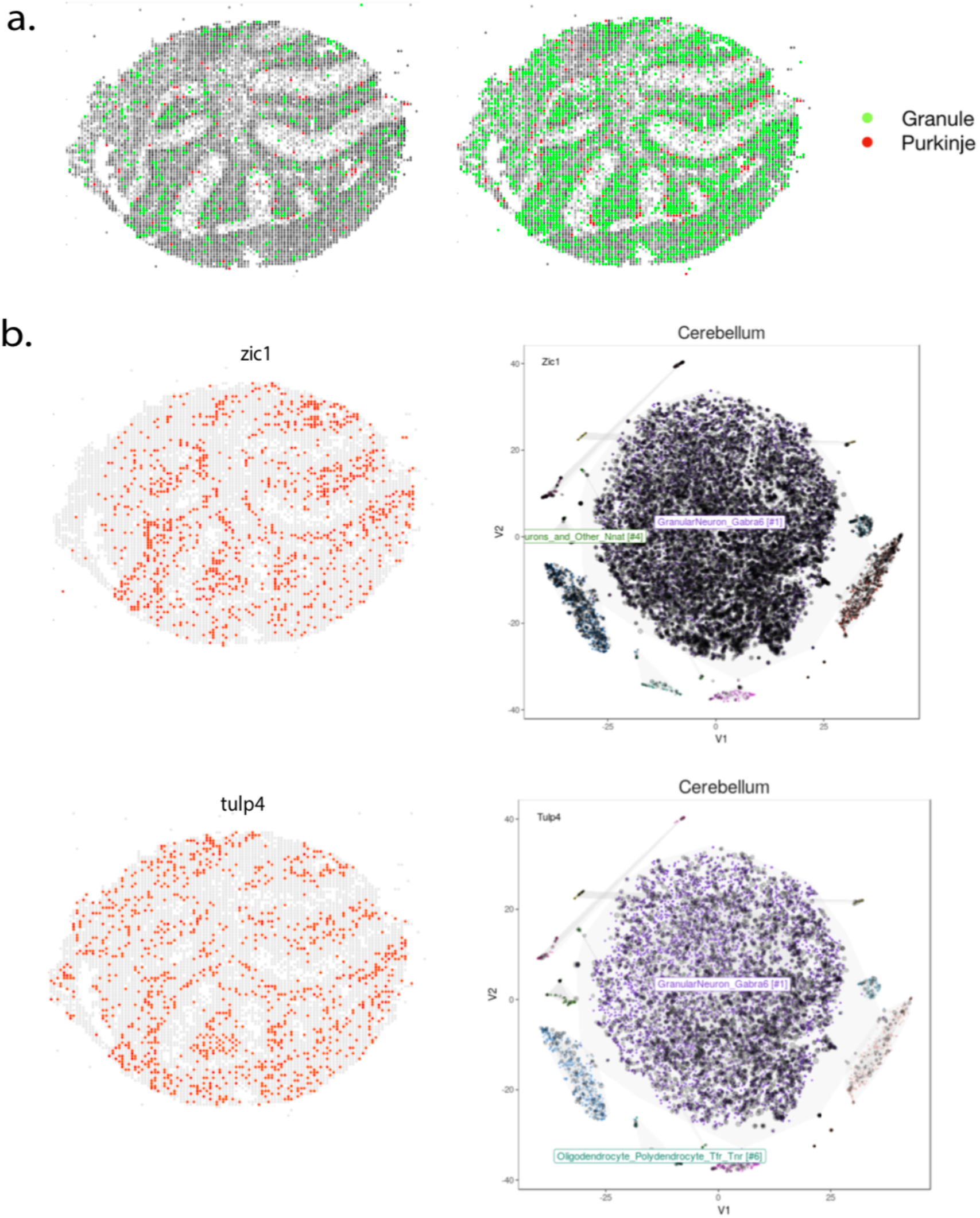
GlmSMA reconstructed the mouse cerebellum accurately. **a**, Mapping results of 5,138 cells from Drop-viz. Reference atlas was established using the Slide-seq data. Most of the granule cells were set to the cerebellar cortex, surrounded by the Purkinje cells. **b**, Reconstructed spatial expression patterns of marker genes. *Zic1* was highly expressed in the granule cells and *tulp4* was highly expressed in the Purkinje cells based on the t-sne plot of the scRNA-seq data. Our reconstructed patterns were consistent with the results from Drop-viz database.

To pinpoint how well glmSMA can work with large numbers of cells in complex tissues, we selected 5,138 mouse-cerebellum cells (4,103 granule, 310 Purkinje) from Drop-Viz. As expected, most of the granule and Purkinje cells were assigned to the correct anatomical structures (Fig. 5a). Per the biological literature, *zic1* and *tulp4* were highly expressed in the granule cells^16^ (Fig. 5b). Our reconstructed patterns were consistent with the results, where the two marker genes were both highly expressed in the granule regions and scarcely expressed in the Purkinje regions. Thus, our mapping results were still robust when we increased the cell number to 5,138.

## GlmSMA accurately reconstructs the mouse hippocampus *in silico* and *in vivo*

We selected mouse-hippocampus tissue to assess glmSMA’s performance in complex tissues with fewer marker genes. Div-seq provided transcripts of 1,188 individual cells with 783 marker genes, and we were able to classify single cells into four cell-types using the pipeline established by Habib et al.^17^ because most were Dente Gyrus (DG) or Cornu Ammonis (CA) cells. We then established the mouse-hippocampus reference atlas from Slide-seq data. One of the mouse-hippocampus’s coronal sections in Slide-seq contained 25,393 locations with 18,896 genes. After selecting the common genes in these two datasets and filtering them by expression levels, 79 marker genes were used for mapping (Extended Data Fig. 10,11,12).

To validate our mapping results, we first investigated the spatial cell types. Most of the DG cells could be correctly assigned in the DG regions. But due to the lack of CA1-specific marker genes that overlap between the two platforms, we falsely mapped several CA1 cells to the DG regions (Fig. 6a). We might solve this problem by adding more CA1-specific genes in the future (Fig. 6c). To further establish that DG cells are region-specific, we combined the hippocampus and cerebellum sections. None of the DG cells were mapped to the cerebellum region. We could not assign previously simulated cerebellum cells to the DG region either. We also investigated well-known DG-specific genes’ reconstructed spatial-expression patterns and found, for example, that tox3 and *pdzd2* were highly expressed in the DG regions. We showed that the reconstructed patterns were consistent with the Hipposeq data^13^ (Extended Data Fig. 16). Taken together, this demonstrates that cell assignments can be verified by marker genes’ spatial cell-types and reconstructed patterns.

**Fig. 6:**
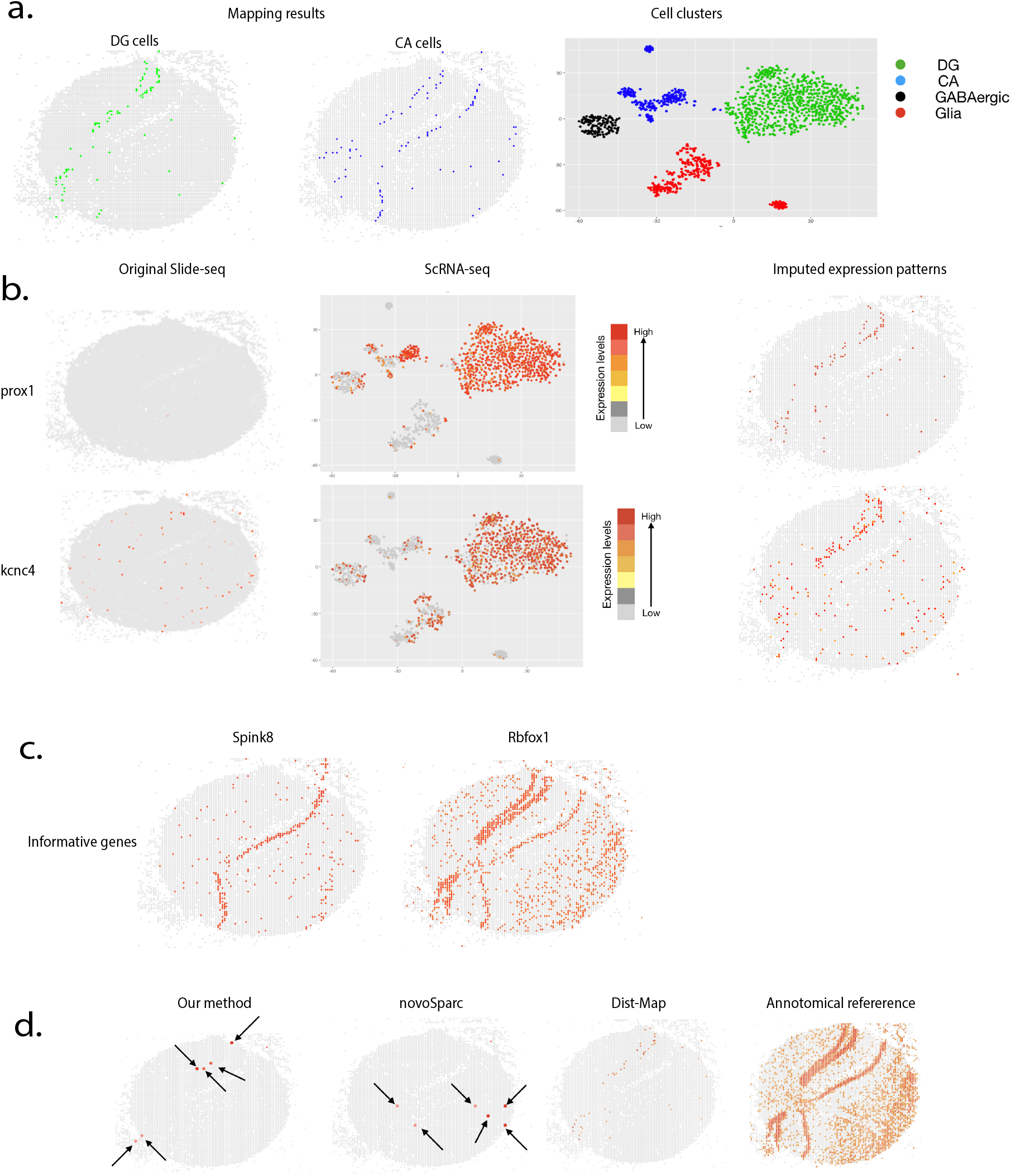
GlmSMA reconstructed the mouse hippocampus. **a**, Mapping results of 1,188 single cells from div-seq. Reference atlas was established by the Slide-seq data. 1,188 cells can be classified as four clusters using predefined marker genes. Most DG cells can be correctly assigned to the DG regions by glmSMA. Some of the CA1 cells were assigned to the DG regions due to the lack the marker genes in the Slide-seq data. The other CA1 cells can be correctly located in the CA1 regions. **b**, Imputed patterns using the mapping results and scRNA-seq profiles. The original Slide-seq cannot capture the genetic patterns of prox1 and kcnc4. GlmSMA imputed the patterns based on the cell locations and their corresponding profiles. The imputed patterns were consistent with the scRNA-seq data. **c**, Informative genes can increase the mapping accuracy. **d**, Comparison between glmSMA, Dist-Map algorithm and novoSparc. Instead of mapping one cell into one location or small patches, Dist-Map assigned one cell into a larger region with close probability. GlmSMA can correctly map one cell into small patches.

We realized that, by combining scRNA-seq and slide-seq, we can impute specific genes’ missing expression profiles into the Slide-seq data and filled the assigned locations with scRNA-seq data from div-seq^17^. *Prox1* and *kcnc4* are well-known marker genes that are only highly expressed in the DG region (Extended Data Fig. 17). The original slide-seq technique cannot capture these two genes’ spatial patterns. By setting the assigned cells with scRNA-seq profiles, the imputed patterns were consistent with the marker genes’ t-SNE plot and Hipposeq (Fig. 6b). Therefore, we can use mapping results and scRNA profiles to successfully impute spatial-expression patterns that the Slide-seq did not capture.

To compare existing algorithms’ cell-assignment prediction accuracy, we used a manually selected marker-gene set to perform glmSMA, novoSparc, and Dist-map in a mouse hippocampus. Fifty marker genes remained after normalization and filtering. We used thresholding to binarize slide-seq and scRNA-seq data expression-profiles as the input for the Dist-map algorithm^2^. Instead of locating cells into limited locations, Dist-map mapped the cells back to large domains. NovoSparc can locate cells into small patches, but assigned many cells to false regions^1^; for example, many DG cells were falsely assigned to the CA1 domain. Compared with previous algorithms, our algorithm can correctly locate most DG cells into small patches in the DG domain, although all three algorithms failed to locate most of the CA1 cells to CA1 regions due to the limited number of CA1-specific genes (Fig. 6d). Thus, our algorithm achieved better performance in mapping cells back to their original locations in the mouse hippocampus.

## Discussion

Here, we proposed a new method to map individual cells back to their original locations that combines reference-atlas and scRNA-seq data. Using both techniques helps us to most accurately measure gene-expression levels in different tissues. ScRNA-seq can provide comprehensive expression profiles for a large number of individual cells, but not location information because of experimental procedure. FISH-based technology, on the other hand, can capture expression profiles at the cellular level but only for a limited number of genes. And although we can use Slide-seq to measure the whole genome in desired tissue simultaneously, only spare genes remain for downstream analysis after normalization and filtering.

Existing single-cell assignment methods have significant limitations. For example, novoSparc only focuses on one type of data. And de novo novoSparc tested their algorithm almost exclusively using the simulated dataset^1^. Other methods, like Dist-map, oversimplify mapping by binarizing the expression profiles in both scRNA-seq data and the reference atlas. While it can work in relatively simple tissues, its performance plummets when applied to more complex tissues like the mouse hippocampus^2^. Our method takes full advantage of the relationship between cell-expressions and locations. With sufficient marker genes, we can locate most cells into correct small patches. Although we cannot locate each individual cell into one unique location in some complex tissues, we can still accurately assign them to relatively small regions.

Other established methods for mapping single-cell data into specific locations are limited to low resolution. For example, MIA and HMRF both focus on locating cell types into regions^9,10^. So, instead of mapping thousands of cells back to several positions, they only map tens of cell-types into large domains. And it is impossible to capture information on individual cells post-mapping because MIA and HMRF merged the expression profiles in the same cell-type. While neither of these methods can locate cells back to small patches, our algorithm assign cells into domains as demonstrated in the intestinal-villus dataset, in which we successfully mapped cells back to 6 large domains with high accuracy. Our algorithm, therefore, provided accurate and high-resolution scRNA-seq data mapping compared with other low-resolution methods.

Our method of mapping individual cells back to their original locations also has limitations. First, our algorithm’s performance relied heavily on marker-gene-set selection. Because building using FISH-like technology to build a reference atlas is so expensive, obtaining enough gene candidates may be difficult^28^. We could overcome this, however, by preprocessing the scRNA-seq or bulk-RNA seq data, which would help to shrink the number of marker gene candidates. Second, current reference-atlas-generating technology, like Slide-Seq, still has significant limitations^6,7^, including poor data-quality needed that prevented us from capturing well-known marker-gene lots’ expression profiles. Despite these challenges, our method still integrated the scRNA-seq data and reference atlas very well, and allowed us to correctly map most of the individual cells to several unique locations.

When combined with the strengths from both scRNA-seq and Spatial-omics techniques, our spatial-expression-pattern analysis and reconstruction identified specific cell organizations in physical space based on the assumption that cells in closer physical proximity are more likely to have similar expression-profiles. ISH images from Allen Brain Atlas and Hipposeq have confirmed reconstructed specific-marker-gene spatial patterns in the mouse hippocampus. Our mapped results also provided an alternative way to impute low-quality data into Slide-seq.

### Method overview

Suppose that the reference atlas has n positions with p genes, and that the scRNA-seq dataset has m cells with the same number of p genes (usually n > m). We aimed to use linear regression to assign the m cells into n positions with L1 and generalized L2 norms via graph Laplacian. First, we created the position-to-gene-expression matrix. Then, based on the matrix, we created one position-to-position graph. If the Euclidean distance between two positions was under a specific threshold, the two positions were connected in the position-to-position graph. We then created a random-walk-normalized-graph Laplacian matrix, which encourage smoothness on coefficients that are connected in the graph.

Our model uses a linear method to measure the differences in gene-expression levels when assigning cells to locations. The optimal solution minimizes the differences between individual cell’s and locations’ gene-expression levels. For each individual cell, we want to minimize the following objective function,

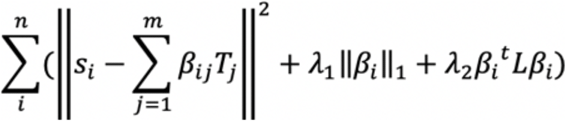

where *s* ∊ *R^n×g^* is the single-cell-expression matrix; *T* ∊ *R^m×g^* is the reference atlas’s marker-gene-expression matrix; L is the normalized graph-laplacian computed from the location-distance graph. The L1 norm-penalization encourages coefficient sparsity, which guarantees that one cell can only be assigned into a few locations. The generalized L2 norm encourages coefficient smoothness, which encourages that cells with similar gene-expression levels are more likely to be assigned into closer locations and that one-cell assignment is more likely to form a patch that scattered distribution. Overall, our method takes full advantage of the relationship between cell-to-cell physical distance and cell-to-location expression distance.

### Data preprocessing

#### Drosophila embryo reference atlas

The embryo consisted of ~6,000 cells and 84 marker genes. Due to the small number of markers genes, we merged 6,000 cells into 3,000 cells by binning them similar to the strategy Karaiskos et al. used with this dataset. We aimed to assign 1,297 cells into 3,039 locations. 1,297 cells can be classified into four cell-types using 84 marker genes.

#### Slide-Seq data normalization

We normalized the data in log-space following the previous pipeline Nitzan et al. Let *d_ij_* be the raw count for gene i in cell j; we normalized it as

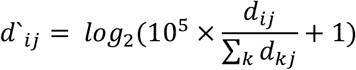

#### Finding highly variable genes in Slide-seq Data and building the reference atlas

For the mouse-hippocampus data, we manually went through all the spatial-expression patterns in the mouse hippocampus to select 79 marker genes and checked the patterns with the Allen Brain Atlas. For the mouse cerebellum data, we manually selected 284 marker genes. We based our reference-atlas construction for different tissues on the marker genes.

### Down-sampling in Slide-Seq datasets

In the Slide-seq datasets, we rounded the physical location coordinates to the next integer multiple of 50 and filtered out low-quality locations where positions with fewer than 50 genes expressed were discarded, resulting in 7,724 cells in the mouse-cerebellum section and 7,790 cells in the mouse-hippocampus section. We then used neighboring locations’ transcriptome sums as the new expression-profiles in the rounded locations.

### Finding highly variable genes in scRNA-seq data

We first normalized and scaled the raw counts in each cell by the median raw-count number across all cells. For each cell, we added a pseudocount of 1 before log transformation. We selected highly variable genes selected based on the Fano factor, defined as 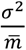 where *σ*^2^ is the gene-expression variance and 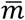 is the expression mean across all cells. We chose a certain mean threshold for the Fano factor.

### Graph Laplacian construction

In physical space, we first constructed an undirected graph G = (V, E), where V is a set of nodes consisting of reference-atlas locations and E is a set of edges consisting of the Euclidean distance between locations. Two locations are connected if the Euclidean distance between them is less than the selected threshold. The random-walk normalized Laplacian matrix on physical locations is defined as,

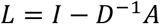

where D is the degree matrix and A is the adjacency matrix in graph G.

The elements of L are shown as,

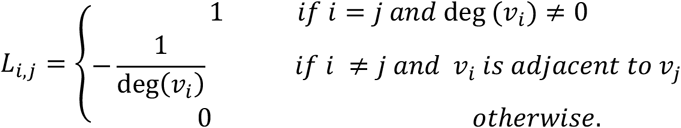

In expression space, we constructed the graph Laplacian similarly. Nodes are cells and edges are the Euclidean distance of expression profiles between two sets of selected marker genes in the new graph G̀.

### Spatial mapping algorithm

To reconstruct spatial expression patterns, our algorithm performs the following steps:

1. Read the gene-expression matrix from scRNA-seq and location-matrix from the reference atlas.
2. Construct two Laplacian matrices.
3. Use CVX to solve for convex function with L1 norm and generalized L2 norm.
4. Assign cells into targeting locations based on the distribution of β in the objective function.
5. Reconstruct the spatial patterns based on the expression-profiles in the scRNA-seq data and cell-locations from the mapped results.

